# A Bayesian account of generalist and specialist formation under the Active Inference framework

**DOI:** 10.1101/644807

**Authors:** Anthony Guanxun Chen, David Benrimoh, Thomas Parr, Karl J. Friston

## Abstract

This paper offers a formal account of policy learning, or habitual behavioural optimisation, under the framework of Active Inference. In this setting, habit formation becomes an autodidactic, experience-dependent process, based upon what the agent sees itself doing. We focus on the effect of environmental volatility on habit formation by simulating artificial agents operating in a partially observable Markov decision process. Specifically, we used a ‘two-step’ maze paradigm, in which the agent has to decide whether to go left or right to secure a reward. We observe that in volatile environments with numerous reward locations, the agents learn to adopt a generalist strategy, never forming a strong habitual behaviour for any preferred maze direction. Conversely, in conservative or static environments, agents adopt a specialist strategy; forming strong preferences for policies that result in approach to a small number of previously-observed reward locations. The pros and cons of the two strategies are tested and discussed. In general, specialization offers greater benefits, but only when contingencies are conserved over time. We consider the implications of this formal (Active Inference) account of policy learning for understanding the relationship between specialisation and habit formation.

**Author Summary:** Active inference is a theoretical framework that formalizes the behaviour of any organism in terms of a single imperative – to minimize surprise. Starting from this principle, we can construct simulations of simple “agents” (artificial organisms) that show the ability to infer causal relationships and learn. Here, we expand upon currently-existing implementations of Active Inference by enabling synthetic agents to optimise the space of behavioural policies that they can pursue. Our results show that by adapting the probabilities of certain action sequences (which may correspond biologically to the phenomenon of synaptic plasticity), and by rejecting improbable sequences (synaptic pruning), the agents can begin to form habits. Furthermore, we have shown our agent’s habit formation to be environment-dependent. Some agents become specialised to a constant environment, while other adopt a more general strategy, each with sensible pros and cons. This work has potential applications in computational psychiatry, including in behavioural phenotyping to better understand disorders.

## Introduction

Any self-organizing system must adapt to its surroundings if it is to continue existing. On a broad timescale, population characteristics change to better fit the ecological niche, resulting in evolution and speciation (1). On a shorter timescale, organisms adapt to better exploit their environment through the process of learning. The degree or rate of adaptation is also important. Depending on the environment around the organism, specialization into a specific niche or favouring a more generalist approach can offer distinct advantages and pitfalls (2). While adopting a single, automatic, behavioural strategy might be optimal for static environments – in which contingencies are conserved – creatures that find themselves in more variable or volatile environments should entertain a broader repertoire of plausible behaviours.

We focus upon adaptation on the shorter timescale in this paper, addressing the issue of behavioural specialisation formally within a Markov decision process formulation of Active Inference (3). Active inference represents a principled framework in which to describe Bayes optimal behaviour. It depends upon the notion that creatures use an internal (generative) model to explain sensory data, and that this model incorporates beliefs about ‘how I will behave’. Under Active Inference, learning describes the optimisation of model parameters – updating one’s generative model of the world such that one acts in a more advantageous way in a given environment (4). Existing work has focussed upon how agents learn the (probabilistic) causal relationships between hidden states of the world that cause sensations which are sampled (4–8). In this paper, we extend this formalism to consider learning of policies.

While it is clear that well-functioning agents can update their understanding of the meaning of cues around them – in order to adaptively modulate their behaviour – it is also clear that agents can form habitual behaviours. For example, in goal-directed versus habitual accounts of decision making (9), agents can either employ an automatic response (e.g. go left because the reward is always on the left) or plan ahead using a model of the world. Habitual responses are less computationally costly than goal-oriented responses; making it desirable to trust habits when they have been historically beneficial (10,11). This would explain the effect of practice – as we gain expertise in a given task, the time it takes to complete that task and the subjective experience of planning during the task diminishes, likely because we have learned enough about the structure of the task to discern and learn appropriate habits (12).

How may our Active Inference agent learn and select habitual behaviours? To answer this question, we introduce a novel feature to the Active Inference framework; namely, the ability to update one’s policy space. Technically, a prior probability is specified over a set of plausible policies, each of which represents a sequence of actions through time. Policy learning is the optimisation of this probability distribution, and optimising the structure of this distribution (i.e. ‘structure learning’) through Bayesian model comparison. Habitual behaviour may emerge through pruning implausible policies, and reducing the number of behaviours that an agent may engage in. If an agent can account for its behaviour without calling on a given policy, it can be pruned, resulting in a reduced policy space, allowing agents to infer which policy it is pursuing more efficiently. Note that in Active Inference, agents have to infer the policy they are pursuing, where this inference is heavily biased by prior beliefs and preferences about the ultimate outcomes. We argue that pruning of redundant behavioural options can account for the phenomenon of specialization (behaviour highly adapted to specific environments), and the accompanying loss of flexibility. In addition to introducing Bayesian model reduction for prior beliefs about policies, we consider its biological plausibility, and its relationship with processes like sleep that have been associated with structure learning (i.e., the removal of redundant model parameters). Finally, through the use of illustrative simulations, we show how optimising model structure leads to useful policies, the adaption of an agent to its environment, the effect of the environment on learning and the costs and benefits of specialization. In what follows, we will briefly review the tenets of Active Inference, describe our simulation set up and then review the behavioural phenomenology in light of the questions posed above.

## Materials and Methods

### Active Inference

Under Active Inference, agents act to minimize their variational free energy (13) and select actions that minimises variational free energy expected following the action. This imperative formalises the notion that an adaptive agent should act to avoid being in surprising states, should they wish to continue their existence. In this setting, free energy acts as an upper bound on surprise and expected free energy stands in for expected surprise or uncertainty. As an intuitive example, a human sitting comfortably at home should not expect to see an intruder in her kitchen, as this represents a challenge to her continued existence; as such, she will act to ensure that outcomes (i.e. whether or not an intruder is present) match her prior preferences (not being in the presence of an intruder); for example, by locking the door.

More formally, surprise is defined as the negative log probability of observed outcomes under the agent’s internal model of the world, where outcomes are generated by hidden states (which the agents have no direct access to, but which cause the outcomes) that depend on the policies which the agent pursues (14):

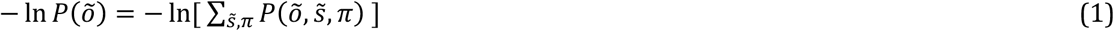

Here, 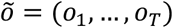 and 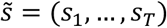 correspond to outcomes (observations) and states throughout time, respectively, and π represents the policies (sequence of actions through time). Since the summation above is typically intractable, we can instead use free energy as an upper bound on surprise (3):

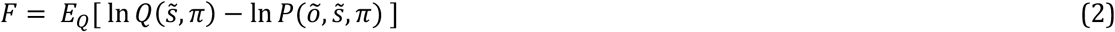

As an agent acts to minimize their free energy, they must also look forward in time and pursue the policy which they expect would best minimize their free energy. The contribution to the expected free energy from a given time, *G(π, τ)*, is the free energy associated with that time, conditioned on the policy, and averaged with respect to a posterior predictive distribution (15):

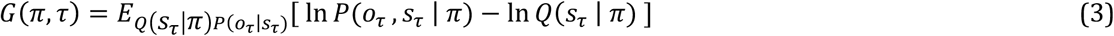

We can then sum over all future time-points (i.e. taking the path integral from the current to the final time: 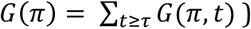 to arrive at the total expected free energy expected under each policy.

### Partially observable Markov decision process and the generative model

A Partially Observable Markov Decision Process (POMDP, or MDP for short) is a generative model for modelling discrete hidden states with probabilistic transitions that depend upon a policy. This framework is useful for formalizing planning and decision making problems and has various applications in artificial intelligence and robotics (16). An MDP comprises two types of

*hidden* variables which the agent must infer: hidden *states* 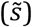 and *policies* (*π*). An MDP agent must then navigate its environment, armed with a generative model that specifies the joint probability distribution of observed outcomes and their hidden causes, and the imperative of minimizing free energy. The states, outcomes and policies are defined more concretely in the following sections.

The MDP implementation consists of the following matrices specifying categorical distributions (6):

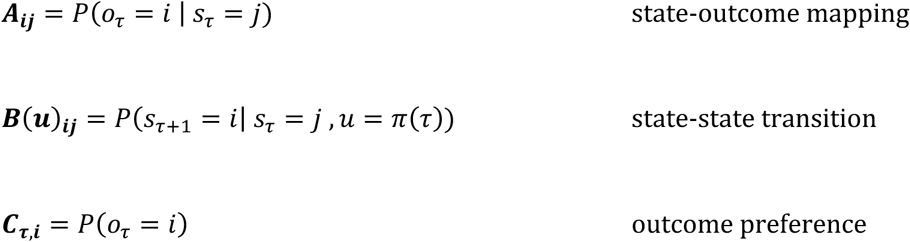

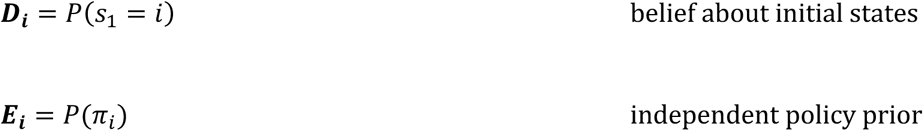

The generative model (Fig 1) assumes that outcomes depend upon states, and that current states depend upon states at the previous timepoint and the action taken (as a result of the policy pursued). Specifically, the state-outcome relationship is captured by an **A** (likelihood) matrix, which maps the conditional probability of any *i-th* outcome given a *j-th* state. A policy, *π_i_* = (*u*_1_, …, *u*_T_), is a sequence of actions (*u*) through time, which the agent can pursue. Generally, an agent is equipped with multiple policies it can pursue. Conceptually, these may be thought of as hypotheses about how to act. As hidden states are inaccessible, the agent must infer its current state from the (inferred) state it was previously in, as well as the policy it is pursuing. State-to-state transitions are described by the **B** (transition) matrix. The **C** matrix encodes prior beliefs about (i.e. a probability distribution over) outcomes, which are synonymous with the agent’s preferences. This is because the agent wishes to minimize surprise and therefore will endeavour to attain outcomes that match the distributions in the **C** matrix. The **D** matrix is the prior belief about the agent’s initial states (the agent’s beliefs about where it starts off). Finally, **E** is a vector of the belief-independent prior over policies (i.e. intrinsic probability of each policy, without considering expected free energy).

**Figure 1:**
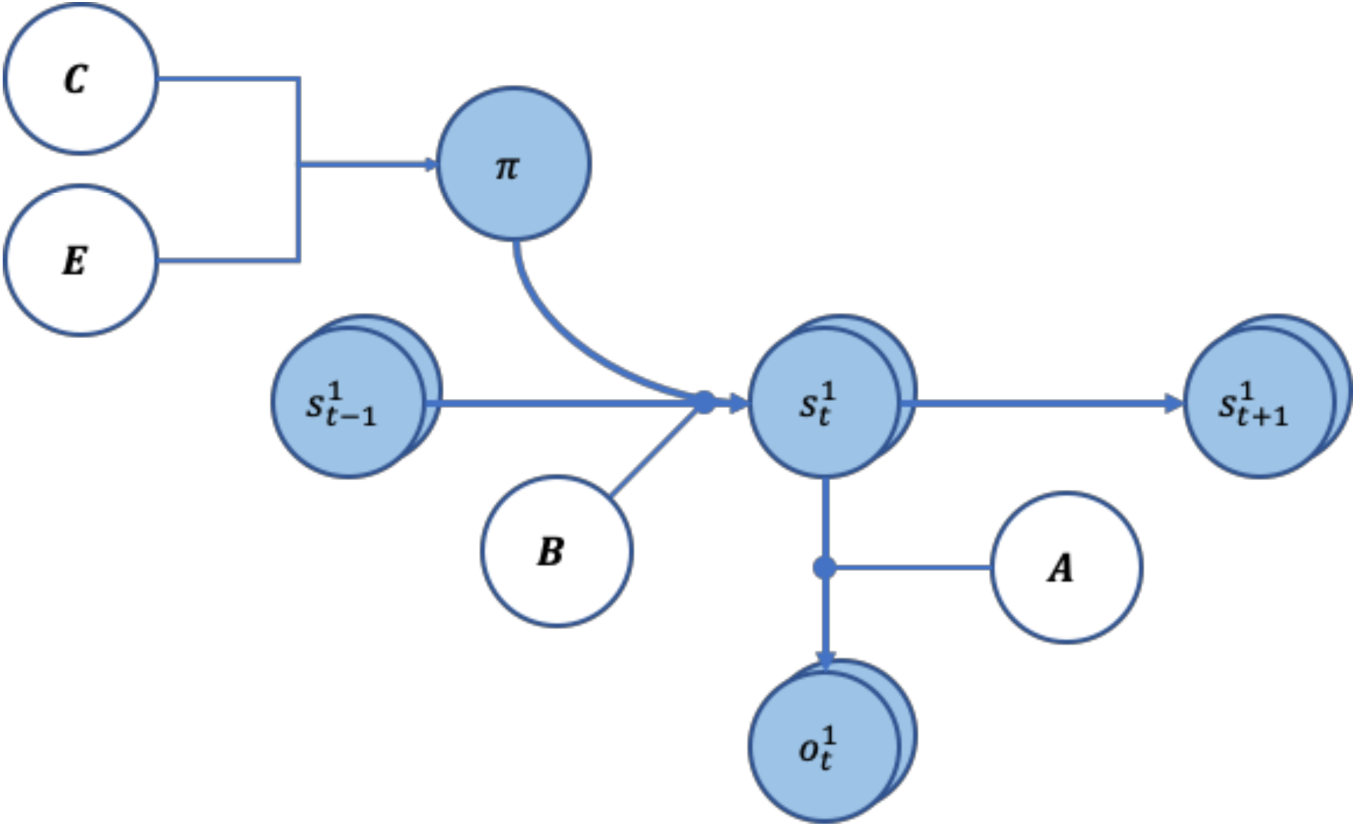
Graphical representation of the generative model. The arrows indicate conditional dependencies, with the endpoint being dependant on where the arrow originated form. The variables in white circles show priors, whereas variables in light blue circles are random variables. The A and B matrices have round arrowhead to show they encode the transition probabilities between the variables.

A concept that will become important below is *ambiguity*. Assuming an agent is in the *i-th* hidden states, *s^i^* the probable outcomes are described by a categorical distribution by the *i-th* column of the **A** matrix. We can therefore imagine a scenario where the distribution *P(O_τ_ | S_τ_ = i)* has *high entropy* (e.g. uniformly distributed), and outcomes are approximately equally likely to be sampled. This is an *ambiguous* outcome. On the other hand, we can have the opposite situation with an *unambiguous* outcome, where the distribution of outcomes given states has *low entropy*. In other words, “if I am in this state, then I will see this and only this”. This unambiguous, precise outcome allows the agent to infer the hidden state that they are in.

Crucially, under Active Inference, an agent must also infer which policy it is pursuing at each time step. This is known as planning as inference (17). The requisite policy inference takes the form:

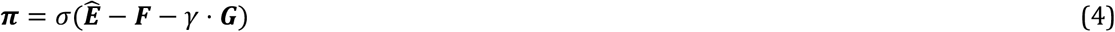

Here, **π** represents a vector of sufficient statistics of the posterior belief about policies: i.e., expectations that each allowable policy is currently in play. ***F*** is the free energy for each policy based on past time points and ***G*** is the expected free energy for future time points. The free energy scores the evidence that each policy is being pursued, while the expected free energy represents the prior belief that each policy will reduce expected surprise or uncertainty in the future. The expected free energy comprises two parts – *risk* and *ambiguity*. Risk is the difference between predicted and preferred outcomes, while ambiguity ensures that policies are chosen to disclose salient information. These two terms can be rearranged into *epistemic* and *pragmatic* components which, as one might guess, reduce uncertainty about hidden states of the world and maximise the probability of preferred outcomes.

The two quantities required to form posterior beliefs about the best policy (i.e., the free energy and expected free energy of each policy) can be computed using the **A**, **B**, and **C** matrices (4,18). The variable γ is an inverse temperature (precision) term capturing confidence in policy selection, and 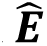 is the (expected log of the) intrinsic prior probabilities in the absence of any inference (this is covered more in-depth in the “*Policy Learning and Dirichlet Parameters*” section below). The three quantities are passed through a softmax function (which normalizes the exponential of the values to sum to one). The result is the posterior expectation; namely, the most likely policy that the agent believes it is in. This expectation enables the agent to select the action that it thinks is most likely.

### Simulations and Task Set-up

We return to our question of the effect of the environment on policy learning via setting up a simulated environment in which our synthetic agent (visualized as a mouse) forages (Figs 2A and 2C). Our environment takes the form of a two-step maze inspired by (19), which is similar to that used in previous work on Active Inference (3,15). The maze allows for an array of possible policies, and the challenge for our agent is to learn to prioritize these appropriately. The agent has two sets of beliefs about the hidden states of the world: where it is in the maze, and where the reward is. The agent also receives two outcomes modalities: *where* it is in the maze and *feedback* received at each location in the maze (Fig 2C, right). The agent always knows exactly where it is in the maze (Fig 2A), and receives different “Feedback” outcomes, depending on where it is in the maze and the location of the reward (Fig 2B).

**Figure 2:**
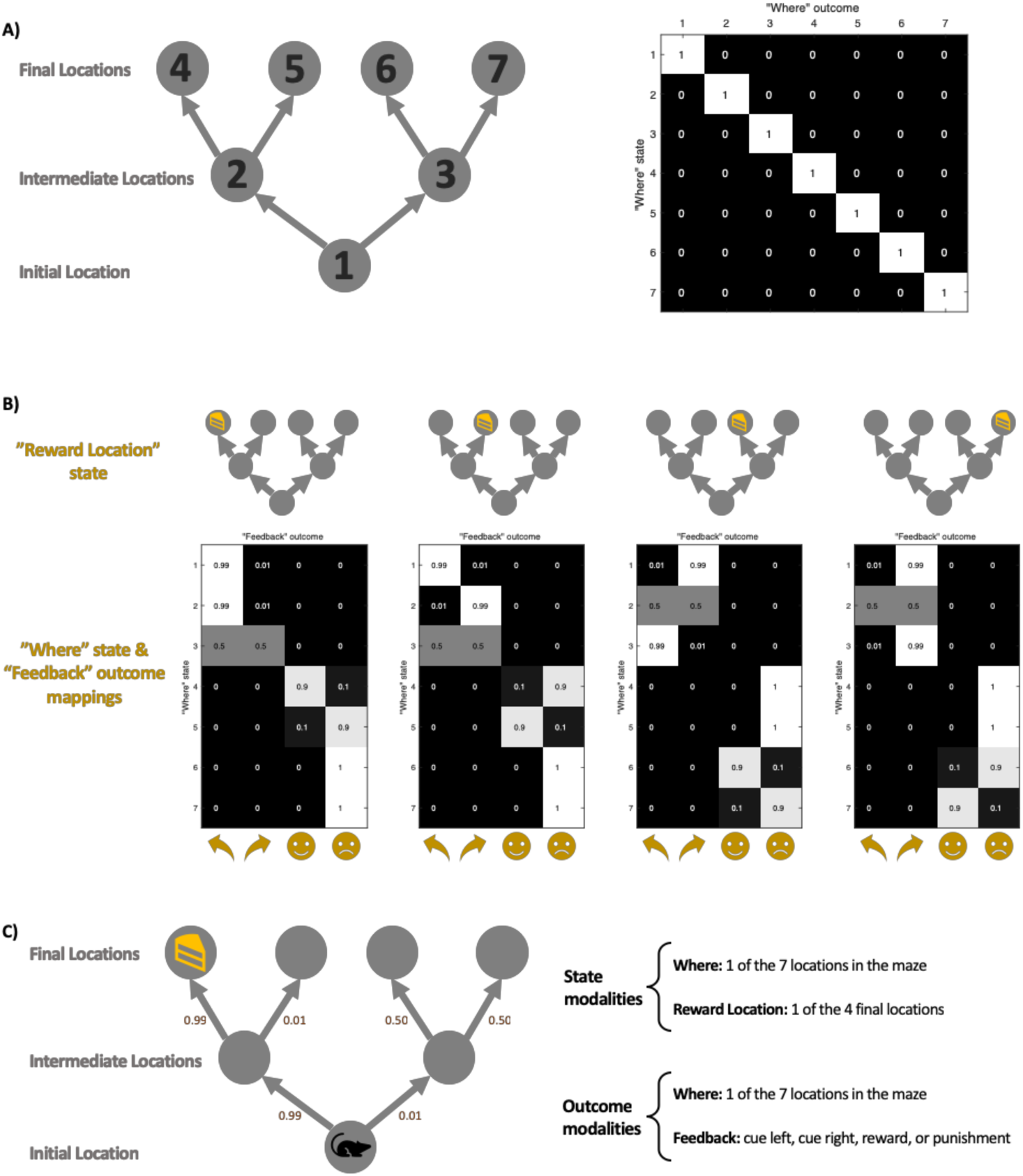
Simulation maze set-up. **(A)** The maze location set-up. There are a total of 7 locations in the maze, each with their corresponding indexes (left diagram). The state-outcome mapping (A matrix) between “Where” (i.e. agent’s current location) state and outcome is an identity matrix (right figure), meaning they always correspond exactly. The maze consists of three stages: initial, intermediate, and final. The state-state transition matrix (B matrix) ensures that an agent can only move forward in the maze, following the direction of the arrow. **(B)** The state-outcome transition probability between the “Where” state and “Feedback” outcome (as encoded by the A matrix). Depending on the location of the reward, the agent receives different feedbacks which include a directional cue (cue left or cue right) in the *initial* and *intermediate locations*, and a reward or punishment at the *final locations*. The index of the y-axis corresponds with the location index in Fig 2A. Here we have depicted *unambiguous cues*, where the agent is 99% sure it sees the cue pointed in the correct (i.e. towards the reward location) cue. **(C)** An example maze set-up with a reward at the left-most *final location*. The agent starts in the *initial location*, and the agent’s model-based brain contains representations of where it is in the maze, as well as where it thinks the reward is. The agent is able to make geographical observations to see where it is in the maze (Fig 2A), as well as receive a “feedback” outcome which gives it a cue to go a certain location, or to give it reward / punishment (Fig 2B). The small numbers beside each arrow illustrate the ambiguity of the cues. As am example, we have illustrated the left-most scenario of Fig 2B.

The mouse always starts in the same initial location (Fig 2A, position 1) and is given no prior information about the location of the reward. This is simulated by setting matrix **D** such that the mouse strongly believes that it is in the “initial location” at τ = 1 but with a uniform distribution over the “reward location”. The agent is endowed with a preference for rewarding outcomes and wishes to avoid punishing outcomes (encoded via the **C** matrix). Cues are placed in the initial and intermediate locations (cue left and cue right). While the agent has no preference for the cues *per se*, it can leverage the cue information to make informed decisions about which way to go to receive the reward. In other words, cues offer the opportunity to resolve uncertainty and therefore have salient or epistemic value. Figure 2C shows the reward in the left-most final location, accompanied by an *unambiguous cue* – the agent is 99% sure that “cue left” means that the reward is actually on the left. This leads it to the correct reward location. The nature of the maze is such that the agent cannot move backward; i.e., once it reaches the intermediate location it can no longer return to the initial location. Once the agent gets to the final location, it will receive either a reward (if it is at the reward location) or be punished.

To see the effect of training under different environments, we set up two different maze conditions: a *volatile environment*, in which the reward can appear in any one of the 4 final locations with equal frequencies, and a *non-volatile environment*, where the reward only appears on the two left final locations (Fig 3A). Crucially, this volatility is between-trial, because these contingencies do not change during the course of a trial. The mouse has no explicit beliefs about changes over multiple trials. Two mice with identical initial parameters are trained in these two distinct environments. With our set-up, each mouse can entertain 7 possible policies (Fig 3B). Four of the policies allow the mouse to get to one of the final four locations, whereas three additional policies result in the mouse staying in either the intermediate or initial locations. Finally, both mice are trained for 8 trials per day for 32 days with *unambiguous* cues in the two environments (Fig 3C). Bayesian model reduction (further discussed below) is performed in-between training – to simulate the effect of sleep and boost learning. Note that we set-up the training environment with *unambiguous cues* to allow for efficient learning, while the testing environment always has *ambiguous cues* – akin to explicit curriculums of school education versus the uncertainty of real-life situations.

**Figure 3:**
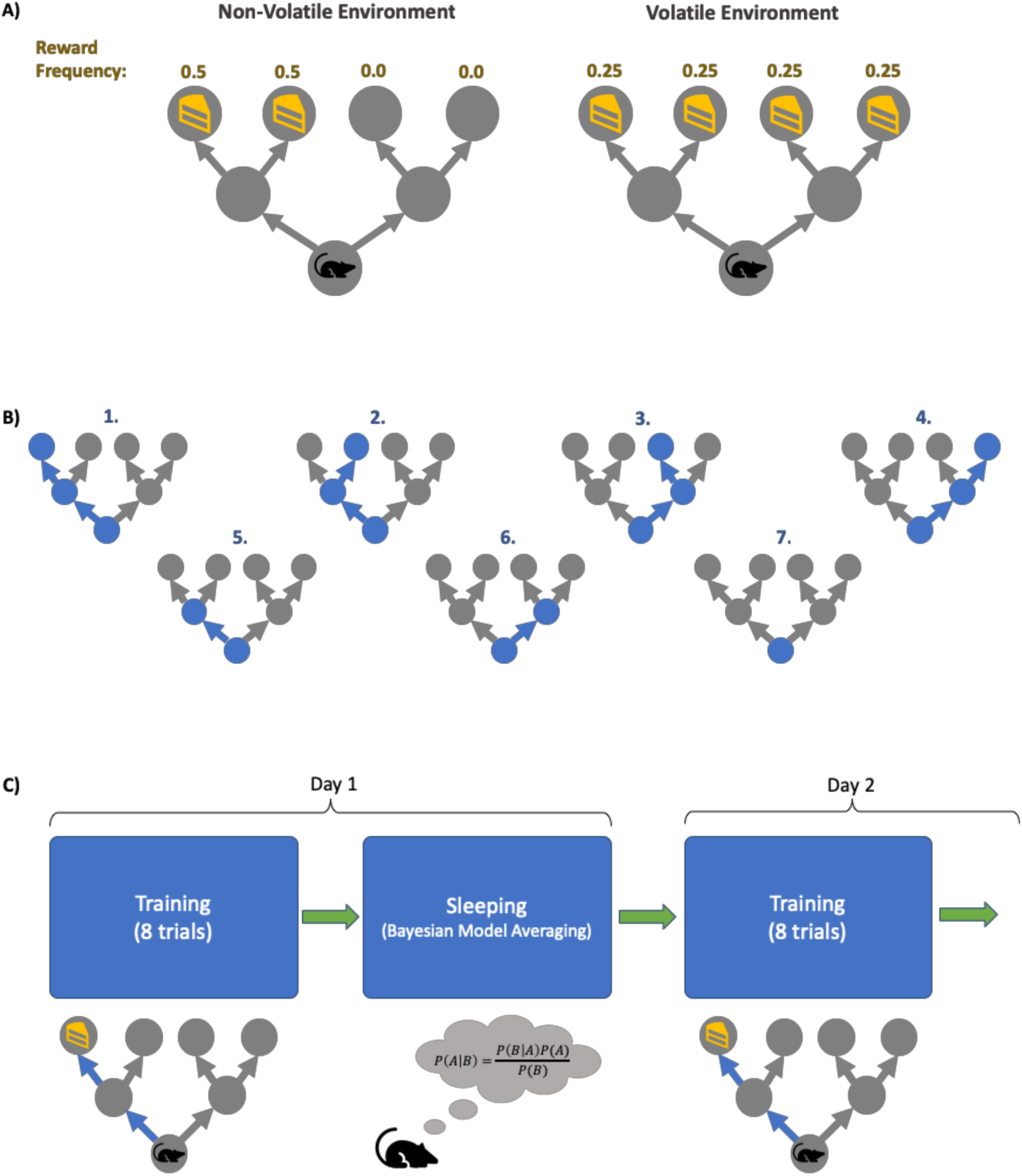
Simulation task set-up. **(A)** The two environments in which the agents are trained. The environment can be *non-volatile* (left), in which the reward always appears on the *left* of the initial location, with equal frequency. The *volatile* environment (right) has reward appearing in all four final locations with equal frequencies. **(B)** The agent’s policies. In our simulation, our agents each have 7 policies it can pursue: the first four policies correspond to the agent going to one of the final locations, policies 5-6 has the agent going to one of the intermediate locations and staying there, and policy 7 has the agent not moving from its initial location for the entire duration of a trial. **(C)** The training cycles. Each day, each agent is trained for 8 trials in their respective environment, and in between days the agent goes to “sleep” (and perform Bayesian model averaging to find more optimal policy concentrations). This process is repeated for 32 days.

### Policy learning and Dirichlet parameters

Whereas inference means optimising expectations about hidden states given the current model parameters, learning is the optimisation of the model parameters themselves (4). Within the MDP implementation of Active Inference, the parameters encode sets of categorical distributions that constitute the probabilistic mappings and prior beliefs denoted by **A**, **B**, **C, D** and **E** above. A Dirichlet prior is placed over these distributions. Since the Dirichlet distribution is the conjugate prior for categorical distributions, we can update our Dirichlet prior with categorical data and arrive at a posterior that is still Dirichlet (20).

While all model parameters can be learned (4,6,20), we focus upon policy learning. The priors are defined as follows:

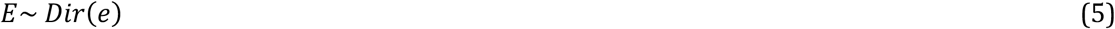

Here *E* is the Dirichlet distributed random variable (or parameter) that determines prior beliefs about policies. The variables *e* = (*e*_1_, …, *e_k_*) are the concentration parameters that parameterise the Dirichlet distribution itself. In the following, *k* is the number of policies. Policy learning occurs via the accumulation of *e* concentration parameters – the agent simply counts and aggregates the number of times it performs each policy and this count makes up the *e* parameters. Concretely, if we define *n_π_* = (*n_π_^1^*, …, *n_π_^k^*) to be the number of times the agent observes itself performing policies *π*^1^, …, *π^k^*, the posterior distribution over the policy space is:

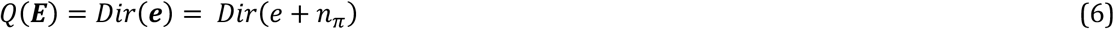

where 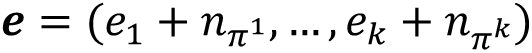 is the posterior concentration parameter. In this way the Dirichlet concentration parameter is often referred to as a “pseudo-count”. Intuitively, the higher the *e* parameter for a given policy, the more likely that policy becomes because more of *Q(**E**)*’s mass becomes concentrated around this policy. Finally, we take the expected logarithm of ***E*** to compute the posterior beliefs about policies in Equation 4:

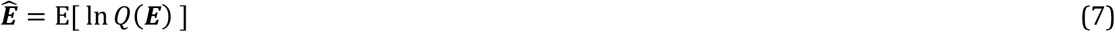

The ***E*** vector can now be thought of as an empirical prior that accumulates the experience of policies that are carried over from previous trials. In short, it enables the agent to learn about the sorts of things that it does. This experience dependent prior policy enters inference via Equation 4. Before demonstrating this experience dependent learning, we look at another form of learning known variously as Bayesian model selection or structure learning.

### Bayesian model comparison

In Bayesian model comparison, multiple competing hypotheses (i.e., models or the priors that defines models) are evaluated in relation to existing data and the model evidence for each is compared (21). Bayesian model averaging (BMA) enables one to use the results of Bayesian model comparison, by taking into account uncertainty about which is the best model. Instead of selecting just the most probable model, BMA allows us to weight models by their relative evidence – to evaluate model parameters that are a weighted average under each model considered. This is especially important in situations where there is no clear winning model (21).

An organism which harbours alternative models of the world needs to consider its own uncertainty about each model. The most obvious example of this is in the evaluation of different plausible courses of action (policies), each entailing a different sequence of transitions. Such models need to be learnt and optimised (22,23) and, rejected, should they fall short. Bayesian model averaging is used implicitly in Active Inference when forming beliefs about hidden states of the world, where each policy is regarded as a model and different posterior beliefs about the trajectory of hidden states under each policy are combined using Bayesian model averaging. However, here, we will be concerned with the Bayesian model averaging over the policies themselves. In other words, the model in this instance becomes the repertoire of policies entertained by an agent.

There is an important connection between these model optimisation procedures, and those processes thought to occur during sleep. This is because a variational free energy minimising creature tries to optimise a generative model that is both *accurate* and *simple* – i.e. that uses the least complicated explanation to describe the greatest number of observations. Mathematically, this follows from the fact that surprise can be expressed mathematically as model evidence – and model evidence is the difference between *accuracy* and *complexity*. During wakefulness, an organism constantly receives perceptual information, and forms accurate yet potentially complex models to explain this (neurobiologically, via increases in the number and strength of synaptic connections through associative plasticity). During sleep, which lacks any precise sensory input, creatures can optimise their models *post hoc* with the goal of reducing complexity (24). This can be achieved by considering *reduced* (simpler) models and seeing how well they explain the data collected during waking hours (22). This is sometimes called Bayesian model reduction and is analogous to the synaptic homeostasis hypothesis of sleep (25). For an excellent review on sleep and model optimisation, see (26), and for a review of Bayesian model reduction, see (27).

Returning to our maze task, our artificial agents traverse through the maze each day and aggregate *e* parameters (Equation 6) to form its daily posterior – that will serve as tomorrow’s empirical prior. During sleep, various reduced models are constructed, via strengthening and weakening amalgamations of *e* parameters. For each configuration of these policy parameters, model evidence is computed and BMA performed to acquire the optimal posterior, which becomes the prior for the subsequent day. In brief, we evaluated the evidence of models in which each policy’s prior concentration parameter was increased by eight, while the remainder were suppressed (by factor of two and four). This creates a model space – over which we can average to obtain the Bayesian model average of concentration parameters in a fast and biologically plausible fashion. Please see S1 Appendix, section A.1 for a general introduction to Bayesian model reduction and averaging. S1 Appendix, section A.2 provides an account of the procedures for an example “day”. In what follows, we now look at the kinds of behaviours that emerge from day-to-day using this form of autodidactic policy learning – and its augmentation with Bayesian model averaging. We will focus on the behaviours that are elicited in the simulations, while the simulation details are provided in the appropriate figure legends (and open access software – see software note).

## Results

### Learning

We now turn to our question about the effect of the environment on policy learning. Intuitively, useful policies should acquire a higher *e* concentration, becoming more likely to be pursued in the future. In simulations, one readily observes that policy learning occurs and is progressive, evident by the increase in *e* concentration for frequently pursued policies (Fig 4), which rapidly reach stable points within 10 days (Fig 4B, see Fig 3C for the concept of “training days”). Interestingly, the relative policy strengths attain stable points at different levels, depending on the environment in which the agent is trained. In a conservative environment, the two useful policies stabilize at high levels (*e* ≈ 32), whereas in a volatile environment, these four useful policies do not reach the same accumulated strengths (*e* ≈ 25). Furthermore, the policies that were infrequently used are maintained at lower levels when trained in a non-volatile environment (*e* ≈ 7), while they are more likely to be considered for the agent trained in the volatile environment (*e* ≈ 11).

**Figure 4:**
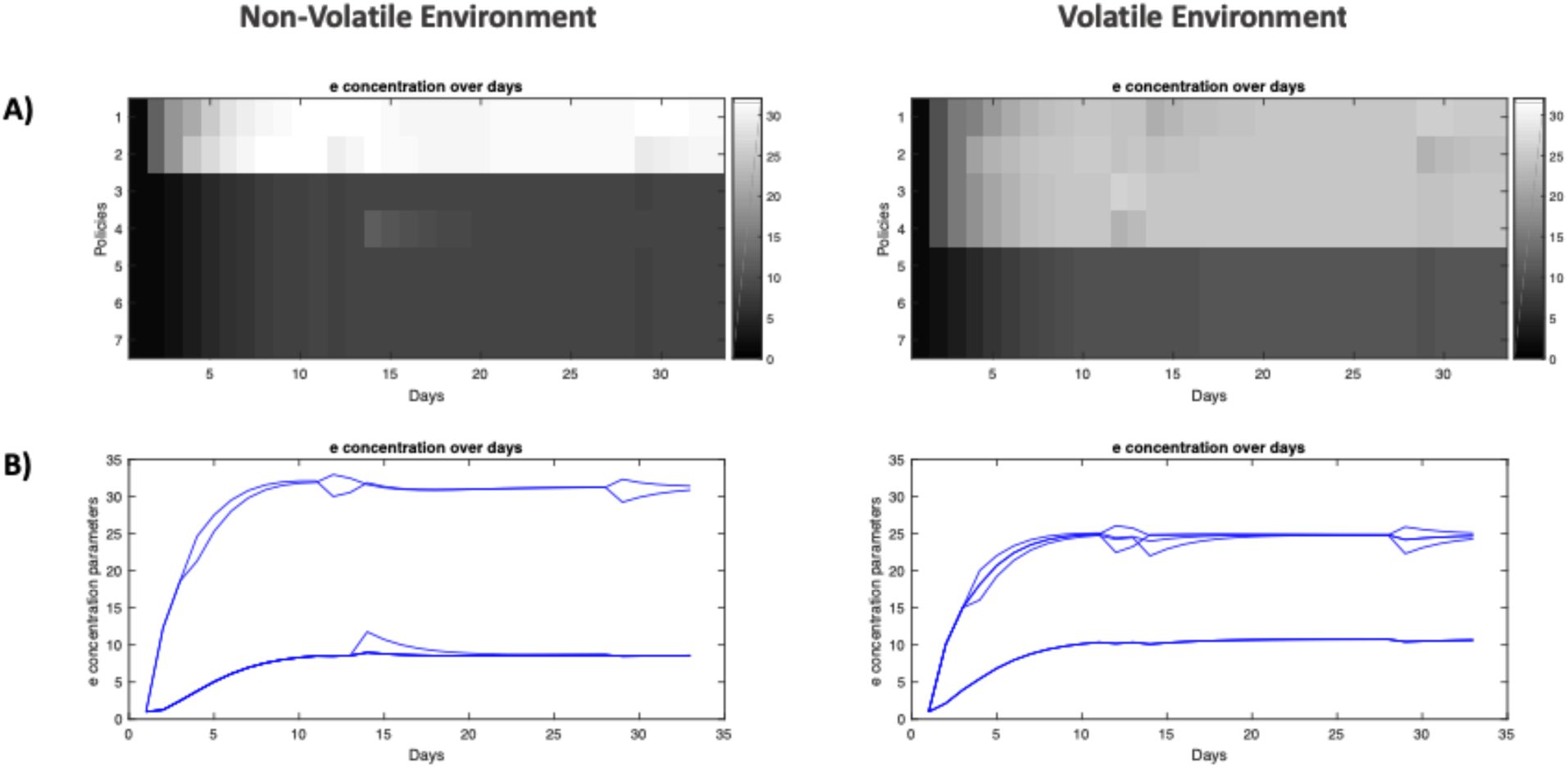
Policy learning over days for agent training in non-volatile and volatile environments. **(A)** Heat-map of *e* concentration parameters for each policy (separated by rows) over all 32 days of training (separated by columns). **(B)** Plot of *e* concentration parameter for policies over 32 days of training.

We will henceforth refer to the agent trained in the non-volatile environment as the *specialist agent*, and the agent trained in the volatile environment as the *generalist agent.* Anthropomorphically, the specialist agent is, *a priori*, more confident about what to do: since the reward has appeared in the leftward location its entire life, it is confident that it will continue to appear in the left, thus it has predilections for left-going policies (policies 1 and 2 of Fig 3B). Conversely, the generalist agent has seen reward appear in multiple locations, thus it experiences a greater level of uncertainty and considers more policies as being useful, even the ones it never uses. We can think of these as being analogous to a general practitioner, who must entertain many possible treatment plans for each patient, compared to a surgeon who is highly skilled at a specific operation.

We can also illustrate the effect of training on the agents’ *reward-acquisition rate:* the rate at which the agents successfully arrive at the reward location (Fig 5). Here, we tested the agents after each day’s training. We see that (Fig 5B, left) with just a few days of training, the specialist agent learns the optimal policies and its *reward-acquisition rate* becomes consistently higher than a *naïve agent* with no preference over any of its policies (*e_native_* = (*e*_1_, …, *e*_7_) = (1, …, 1)). Conversely, the generalist agent never becomes an expert in traversing its environment. While it learns to identify the useful policies (Fig 5A, right), its performance is never significantly better than the naïve agent (Fig 5B, right). Overall, we see that a *non-volatile* environment leads to specialization, whereas a *volatile* environment leads to the agent becoming a generalist.

**Figure 5:**
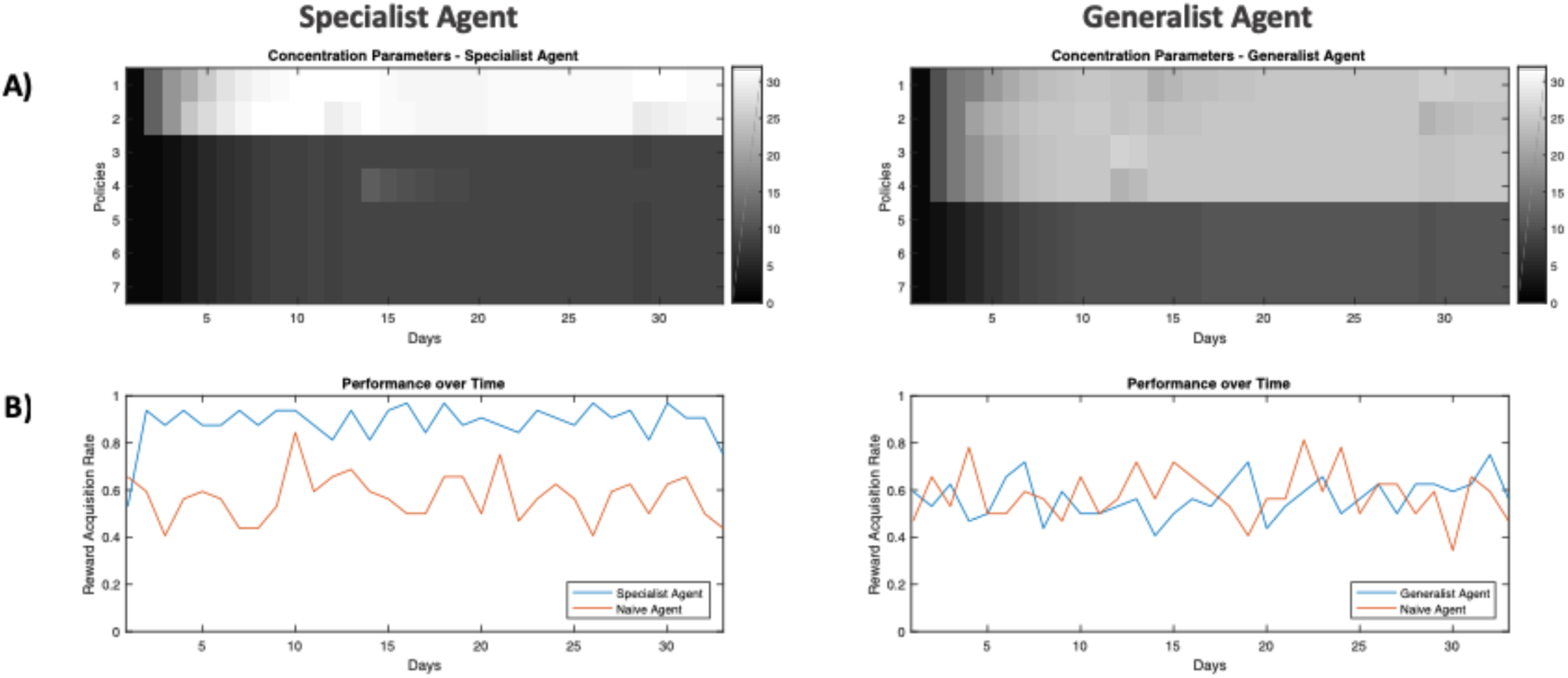
Example performance of in-training agents over days. **(A)** Heat-map of *e* concentration parameters for each policy (separated by rows) over all 32 days of training (separated by columns). **(B)** The frequency at which the agent is able to get to the reward location when tested under ambiguity. This simulated testing is done after each day of training, where each agent is tested under ambiguity (the agent is 65% sure it sees the correct cue) for 32 trials, where the reward location / frequency in the testing environment is identical to the environment in which the agent is trained (i.e. a specialist agent is tested in an environment with low volatility and the reward always being on the left of the initial location). The frequency is computed from how many out of the 32 trials the agent is able to get to the true reward location.

### Testing

We then asked how the specialist and generalist mice perform when transported to different environments. We constructed three testing environments (Fig 6A): the *specialized environment,* similar to the environment the specialized agent is trained in; namely, with rewards that only appear on the left side of the starting location (low volatility); the *general environment* containing rewards that may appear in any of the four final locations (high volatility); additionally, the *novel environment* has reward *only* on the right side of the starting location (low volatility).

**Figure 6:**
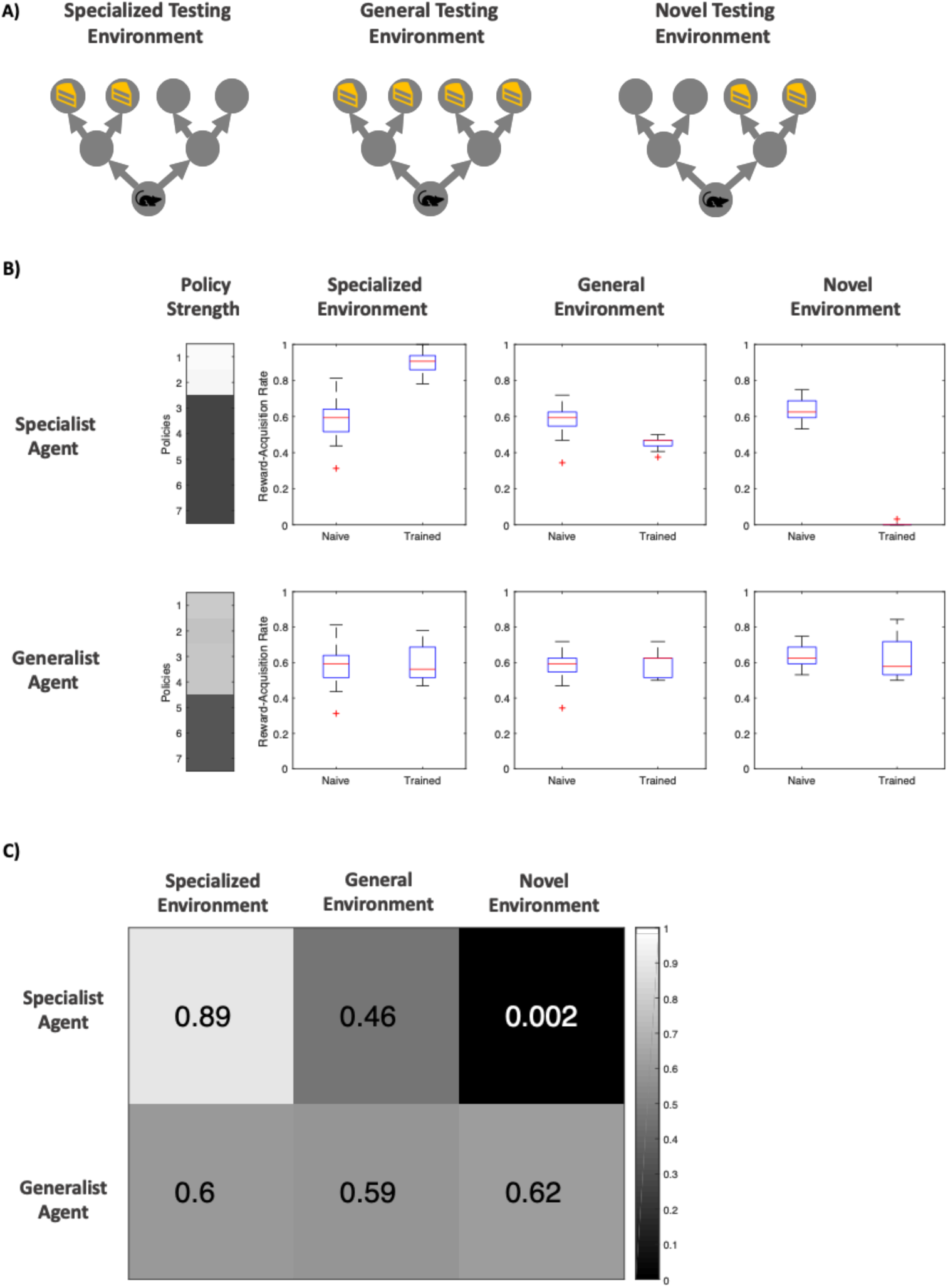
Post-training performance of specialist and generalist agents in ambiguous environments. (the agent is 65% sure it sees the cue telling it to go in the correct direction) **(A)** Visualization of the three testing environments. The *specialized* and *general* testing environment have identical reward location and frequencies to the environments in which the *specialist* and *generalist* agents were trained, respectively. The novel environment is a new, low volatility environment in which the reward only appears to the *right* of the initial location. **(B)** Distribution of reward-acquisition-rate of specialist and generalist agents compared against a naïve agent with no training. The “Policy Strength” column shows how much of each policy the agent has learned, and the three boxes of boxplots show the comparison in performance. The reward-acquisition rate distribution is generated via running each trial 32 times to generate a reward-acquisition rate (proportion of times the agent correctly navigates to the reward location), and repeating this process 16 times to generate a distribution of scores. **(C)** A confusion matrix of mean reward-rate of each agent within each testing environment. Both the heat map and the colour over each element represents the reward-acquisition rate.

Each agent was tested for 512 trials in each test environment. Note that the agents do not learn during the testing phase – we simply reset the parameters in our synthetic agents after each testing trial to generate perfect replications of our test settings. We observe that an untrained (naïve) agent has a baseline reward-acquisition rate of ∼60%. On the contrary, the specialist agent excels when the environment is similar to that it trained in, performing at the highest level (89%) out all the agents. In contrast, the specialist agent performs poorly in a general environment (46% reward-acquisition), and fails all but one out of its 512 attempts in a novel environment where it needs to go in the opposite direction to that of its training (Figs 6B and 6C). The generalist agent, being equally trained in all four policies – that take it to one of the end locations – does not suffer from reduced reward-acquisition when exposed to a new environment (the specialized environment or novel environment). However, it does not perform better in a familiar, general environment either. The agent’s reward-acquisition remains around 60% across all testing environments, similar to that of a naïve agent (Figs 6B and 6C).

Overall, we find that becoming a specialist versus a generalist has sensible trade-offs. The benefit of specialization is substantial when operating within the same environment, consistent with data on this topic in a healthcare setting (28,29). However, if the underlying environment is different, then performances can decrease to one which is poorer than the performance without specialization. Drawing once again from healthcare, the benefits of generalising are numerous as it allows for the practitioner to react more flexibly to changing demography and societal perspectives (30). Conversely, being a generalist means the agent never thrives in a single environment.

## Discussion

### Specialists and Generalists

Our focus in this paper has been on policy optimisation, where discrete policies are optimised through learning and Bayesian model reduction. By simulating the development of specialism and generalism, we illustrated the capacity of a generalist to perform in a novel environment, but its failure to reach the level of performance of a specialist in a specific environment. We now turn to a discussion of the benefits and costs of expertise. Principally, the drive towards specialization (or expertise) is the result of the organism’s imperative to minimize free energy. As free energy is an upper bound on surprise (negative Bayesian model evidence), minimizing free energy maximizes model evidence (31). As model evidence takes into account both the accuracy and complexity of an explanation (22), it is clear that having a parsimonious model that is well-suited to the environment – a specialist model – will tend to minimize free energy over time, provided the environment does not change.

In a stable (conservative, non-volatile) setting, a complex environment can be distilled down into a simple model without sacrificing accuracy. This results in efficient policy selection and provides a theoretical framework for understanding the formation of expertise. In our simulations, the agent trained in the unchanging environment learns to favour the two policies that go left, as the reward is always on the left of the starting location. It thus becomes more efficient and acts optimally in the face of uncertainty. This is evident by its excellent performance in finding left-situated rewards (Fig 6). Indeed, previous theories of expertise differentiate experts from novices in their ability to efficiently generate complex responses to their domain-specific situations (32–34). For example, in typists, expertise is most well-characterized by the ability to quickly type different letters in succession using different hands (33,35). In essence, the expert needs to quickly select from her repertoire of motor policies the most appropriate to type the desired word. This is a non-trivial problem: using just the English alphabet, there are a total of 26*^m^* ways of typing an *m*-character-long word (e.g. a typist needs to select from 26^6^ = 308915776 policies to type the 6-letter word “EXPERT”). It is no wonder that a beginner typist struggles greatly and needs to forage for information by visually searching the keyboard for the next character after each keystroke. The expert, on the other hand, has an optimised prior over her policy space, and thus is able to efficiently select the correct policies to generate the correct character sequences.

However, specialization does not come without its costs. The price of expertise is reduced flexibility when adapting to new environments, especially when the new settings are contradictory to previous settings (11,36). Theoretically, the expert has a simplified model of their domain, and, throughout their extensive training, has the minimum number of parameters necessary to maintain their model’s high accuracy. Consequently, it becomes difficult to fit this model to data in a new, contradictory environment that deviates significantly from the expert’s experience. For instance, we observe that people trained in a perceptual learning task perform well in the same task, but perform worse than naïve subjects when the distractor and target set are reversed – and take much longer to re-learn the optimal response than new subjects who were untrained (37).

Conversely, a volatile environment precludes specialization. The agent cannot single-mindedly pursue mastery in any particular subset of policies, as doing so would come at the cost of reduced accuracy (and an increase in free energy). The generalist agent therefore never reaches the level of performance that the specialist agent is capable of at its best. Instead, the generalist performs barely above the naïve average reward-acquisition rate, even when tested under a general environment. However, the generalist is flexible. When placed in novel and changing environments, it performs much better than our specialist agent.

Interestingly, we note that specialist formation requiring a *conservative* training environment adheres to the requirements specified by K. Anders Ericsson in his theory of *deliberate practice* – a framework for any individual to continuously improve until achieving mastery in a particular field (34,38,39). Ericsson establishes that deliberate practice requires a well-defined goal with clear feedback (low volatility learning environment) and ample opportunity for repetition and refinement of one’s performance (training, repetition and, potentially, Bayesian model reduction during sleep).

### Ways of Learning

There are two principal modes of (policy) learning. The first is *learning via reduction,* which entails a naïve agent that starts with an over-complete repertoire of possible policies, who then learns to discard the policies that are not useful. This is how we have tackled policy learning here; specifically, via optimising a Dirichlet distribution over policies, using Bayesian model reduction. By starting with an abundance of possible policies, we ensure that the best policy is likely to always be present. This also corresponds with the neurobiological findings of childhood peaks in grey matter volume and number of synapses, followed by adolescent decline (40–42). In this conceptualization, as children learn they prune away redundant connections, much as our agents triage away redundant policies. Likewise, as the policy spaces are reduced and made more efficient, we also observe a corresponding adolescent decline in brain glucose usage (43). This is consistent with the idea that informational complexity is metabolically more expensive (44).

The second method of learning is *learning via expansion*. Here, we start with a very simple model and increase its complexity until a more optimal model is reached. Concretely, this problem of increasing a parameter space is one addressed by Bayesian Nonparametric modelling (45), and has been theorized to be utilized biologically for structure learning to infer hidden states and the underlying structures of particular situations (46,47).

### Hyperpriors and Evolution

Note that the way in which we define our reduced model influences how learning of the *e* parameters proceeds. Recall that to explore a plausible model space of priors, we increased concentration parameters by 8 and divided the others by either 2 and 4. These changes were hand-crafted and somewhat arbitrary, and are basically used to assess the change in model evidence when prior beliefs in a particular policy are strengthened, relative to others. The exact ways in which the repertoire of reduced models could be specified in terms of as *hyperparameters,* and reasonably there would be *hyperpriors,* which are prior distributions over hyperparameters.

Similar to model parameters, hyperpriors can be optimised over time to reduce the path integral of free energy. For example, in Bayesian model reduction there can be different settings for how much to increment concentration parameters, and different degrees of comprehensiveness when it comes to exploring the reduced model space (i.e. whether or not to iterate through all possible combinations of policies). If one subscribes to the notion that this kind of structure learning occurs during sleep, optimising hyperparameters becomes a behavioural scheduling problem. The organism can sleep more frequently to compute empirical priors for the next period of waking, or it can spend more time awake to gather empirical data. Both periods (sleep and wake) offer different advantages, and the balance between them is a delicate equilibrium – influenced by ecological pressures. One can imagine that the species-specific circadian rhythms maintain this optimum and evolution helps to fine-tune the hyperparameters facilitating this schedule (48,49).

### Bayesian model comparison

In our simulations, we optimised policy strengths through the process of Bayesian model reduction (to evaluate the free energy or model evidence of each reduced model), followed by model averaging – in which we take the weighted average over *all* reduced models. However, BMA is just one way of using model evidences to form a new model. Here, we discuss other approaches to model comparison, their pros and cons, and biological implications. The first is Bayesian model *selection*, in which only the reduced model with the greatest evidence is selected to be the prior for the future, without consideration of competing models. This offers the advantage of reduced computational cost (no need to take the weighted sum during the averaging process) at the cost of a myopic selection – the uncertainty over reduced models is not taken into account.

The second method, which strikes a balance between BMA and Bayesian model selection with respect to the consideration of uncertainty, is BMA with *Occam’s Window* (50). In short, a threshold is established, *O_R_*, and if the log evidence of any reduced model is not within *O_R_*, we simply do not consider that reduced model. Neurobiologically, this would correspond to the effective silencing of a synapse if it falls below a certain strength (51). This way, multiple reduced models and relative uncertainties are still considered, but a great degree of computational cost is saved since less reduced models are considered overall.

Interestingly, the Occam’s window itself can also be thought of as a hyperprior. A wide window (high *O_R_*) means more models are considered, which offers a more optimal averaged prior but at higher computational costs; a narrow window (low *O_R_*) means only the models with high model evidence are considered. This allows for efficient averaging over only the best models but comes at the cost of strict pruning. Likewise, both strategies offer different advantages, and the optimal balance may depend on the nature of the agent’s environment (i.e., is it an environment that provides definitive evidence for a small number of policies – or is it an ambiguous environment?). A deviation from the optimum may result in reduced fitness and suboptimal inference – a potentially useful perspective on psychopathology in neuropsychiatric illnesses. For instance, an overly strict pruning rule – while being highly efficient for policy optimisation – may result in useful policies being forever lost. This sub-optimal form of structure learning may relate to the aberrant pruning which in schizophrenic patients (52), leading to maladaptive policy spaces and policy-derived priors that could drive hallucinations.

### Limitations

One limitation of our simulations was that our agents did not learn about cues at the same time they were learning about policies; in fact, the agents were constructed with priors on which actions were likely to lead to rewards, given specific cues (that is, a correctly perceived cue-left was believed by the agents to – and actually did – always lead to a reward on the left). As such, we did not model the learning of cue-outcome associations and how these may interact with habit formation. We argue this is a reasonable approximation to real behaviour; where an animal or human first learns how cues are related to outcomes, and, once they have correctly derived a model of environmental contingencies, can then proceed to optimising policy selection.

Additionally, while we were able to see a significant performance difference between specialist and generalist agents, there was little distinction between the performance of generalist and naïve agents. This likely resulted from the “two-step” maze being a relatively simple task. As agents are incentivized to go to the very end of the maze to receive a reward, the naïve agents are not at a disadvantage compared to generalists (since both have equal prior beliefs about the final locations). An alternative explanation is that the generalist strategy is simply the preservation of naivety.

To address the above limitations, future work could involve more complex tasks to more clearly differentiate between specialist, generalist and naïve agents. Additional types of learning should also be included, such as the learning of state-outcome mappings (optimising the model parameters of the likelihood (**A**) matrix, as described in (4,6)), to understand how learning of different contingencies influence one another. In addition, more complex tasks may afford the opportunity to examine the generalisation of specialist knowledge to new domains (53). This topic has recently attracted a great deal of attention from the artificial intelligence community (54,55).

Furthermore, it would be interesting to look at policy learning using a hierarchical generative model, as considered for deep temporal models (56). This likely leads to a more accurate account of expertise-formation, as familiarity with a domain-specific task should occur at multiple-levels of the neural-computation hierarchy (e.g. from lower level “muscle memory” to higher level planning). Likewise, more unique cases of learning can also be explored, such as the ability and flexibility to re-learn different tasks after specializing, the influence of sleep deprivation on policy learning, and different ways of conducting model comparison (as discussed above).

## Conclusion

In conclusion, we have presented a computational model under the theoretical framework of Active Inference that equips an agent with the machinery to learn habitual policies via a prior probability distribution over its policy space. In our simulations, we found that agents who specialize – employing a restricted set of policies because these were adaptive in their training environment – can perform well under ambiguity but only if the environment is similar to its training experiences. On the contrary, a generalist agent can more easily adapt to changing, ambiguous environments, but is never as successful as a specialist agent in a conservative environment. These findings cohere with the previous literature on expertise formation – as well as with common human experience. Finally, these findings may be important in understanding aberrant inference and learning in neuropsychiatric diseases.

## Acknowledgments

Rosetrees Trust (Award Number 173346) to T.P. K.J.F. is a Wellcome Principal Research Fellow (Ref: 088130/Z/09/Z).

## Appendix A: Bayesian model comparison

### A.1 Derivations of Bayesian Model Comparison

We start off with the following approximate equalities of the approximate posterior of a set of parameters (26):

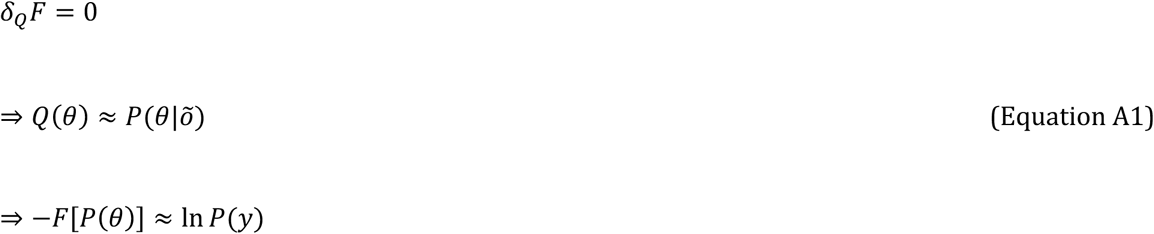

*θ* = (*θ*_1_, *θ*_2_, …) is used here to denote any arbitrary set of parameters, and *δ_Q_F* = 0 means the variation of the free energy with respect to the approximate posterior is zero (i.e. a stationary point of the free energy). For the purpose of policy learning as discussed in this paper, it would be identical to substitute the tuple of concentration parameters, *e*, in lieu of *θ* below.

In order to perform Bayesian model comparison, we define our two models: a *full* model (in this case, the model the agent used in the previous day), and a *reduced* model (model constructed during sleep which the agent compares against the full model). We define the probabilities under the two models with the *P_F_* and *P_R_*, respectively. Crucially, we make a key assumption, that the likelihood of observing the outcomes is equally likely under both models:

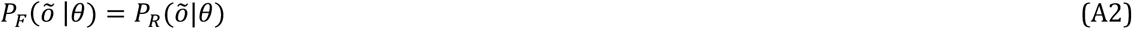

We begin by writing out Bayes rule to both the full and reduced models:

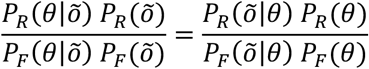

Using the equality in Equation A2 to cancel the likelihood terms, and rearranging, we arrive at the following equality:

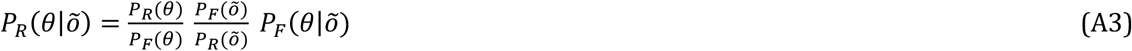

Integrating both sides:

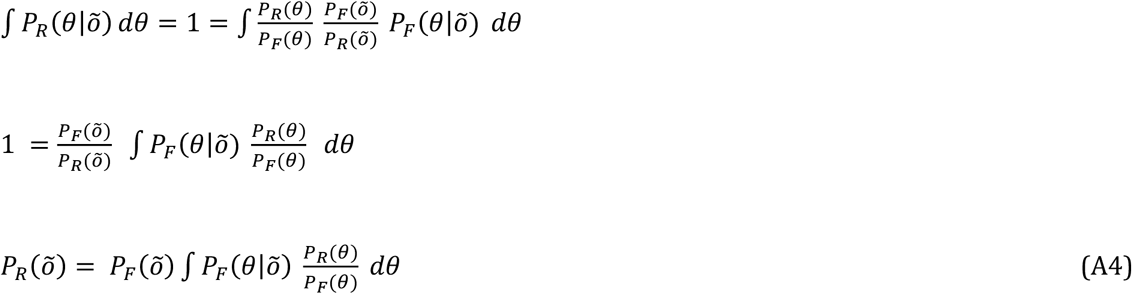

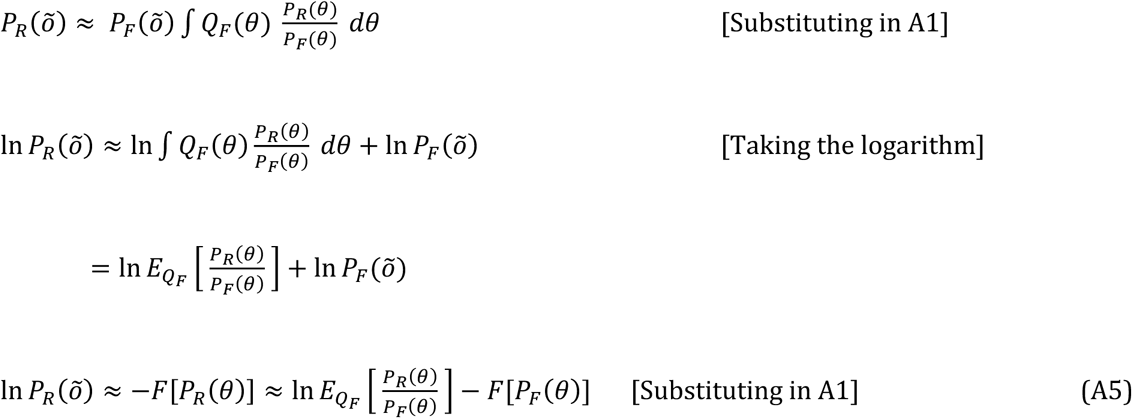

Equation A5 tells us that the model evidence of any reduced model can be evaluated given the prior of the reduced and full models, and the evidence of the full model. Applying the above knowledge to the *e* concentration parameters defined previously, we have the following:

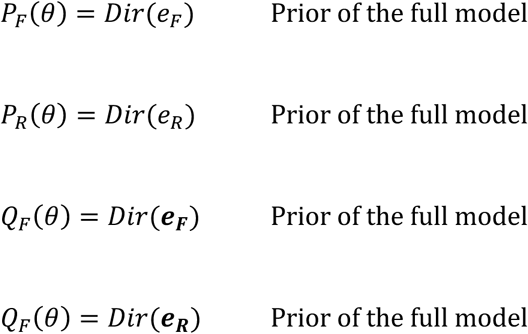

In order to compare relative model evidence, we look at the log ratio of the reduced and full model evidence, which is the same as the difference in their free energy (free energy of the full model minus the reduced):

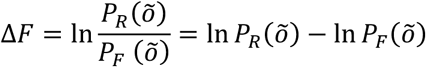

In the discrete case, the above can simply be re-written with Beta functions Β(⋅) (26):

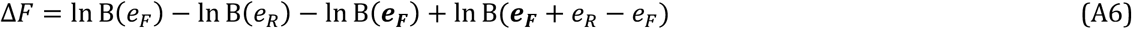

We can apply the above to any reduced model to evaluate its evidence relative to the full model. Intuitively, the higher Δ*F* is, the more evidence the reduced model has. We can evaluate Δ*F* for an arbitrarily large number of reduced models.

In the case of *Bayesian model selection*, the reduced model with the highest model evidence is selected as the optimal model. That is to say, given a vector of the relative free energy for each reduced model, ΔF, we pick the *e_R_* which gives max (ΔF). However, since we are interested in *Bayesian model averaging*, we need to compute the probability of each reduced model within the entire reduced model space we defined:

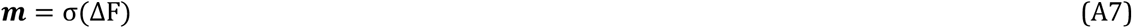

where ***m**_i_* = *Q*(*m* = *i*) is the posterior probability of each reduced model and Σ is the softmax function, 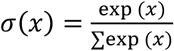, which squashes the set of values in vector ΔF into a range that is between

[0, 1] and sums to 1 (i.e. forms a probability distribution). After the probability of each reduced model is computed, we simply take a weighted sum of each reduced model parameters, weighted by their probability, to get the final, Bayesian model averaged parameters:

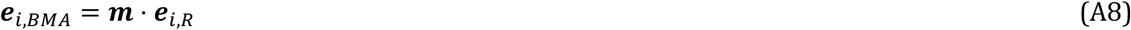

where ***e**_i,R_* is a vector of the *i-th* concentration parameters for each reduced model, and ***e**_i,BMA_* is the *i-th* Bayesian model averaged concentration parameter over all reduced models.

### A.2 Example application of Bayesian model comparison to maze task

Taking our “two-step” maze task for example, let us imagine an agent that repeatedly pursues policy 1 (Fig 3B) throughout the day. At the end of the day, having completed 8 trials, its *e* parameter for policy 1 has increased from a prior concentration of 1 to a posterior concentration of 9 (Fig A1a). The agent then goes to sleep, where it entertains possible combinations of reduced models for prior *e* parameters (Fig A1b) and computes the model evidence for each reduced model using the derivations shown in Appendix section A.1 (the resulting model evidence is shown in Fig A1c). Specifically, it tests different initial parameters (Fig A1b) in lieu of the true prior parameters used (Fig A1a, left) to see whether these provide better explanations for the observed data (Fig A1a, right).

**Figure A1:**
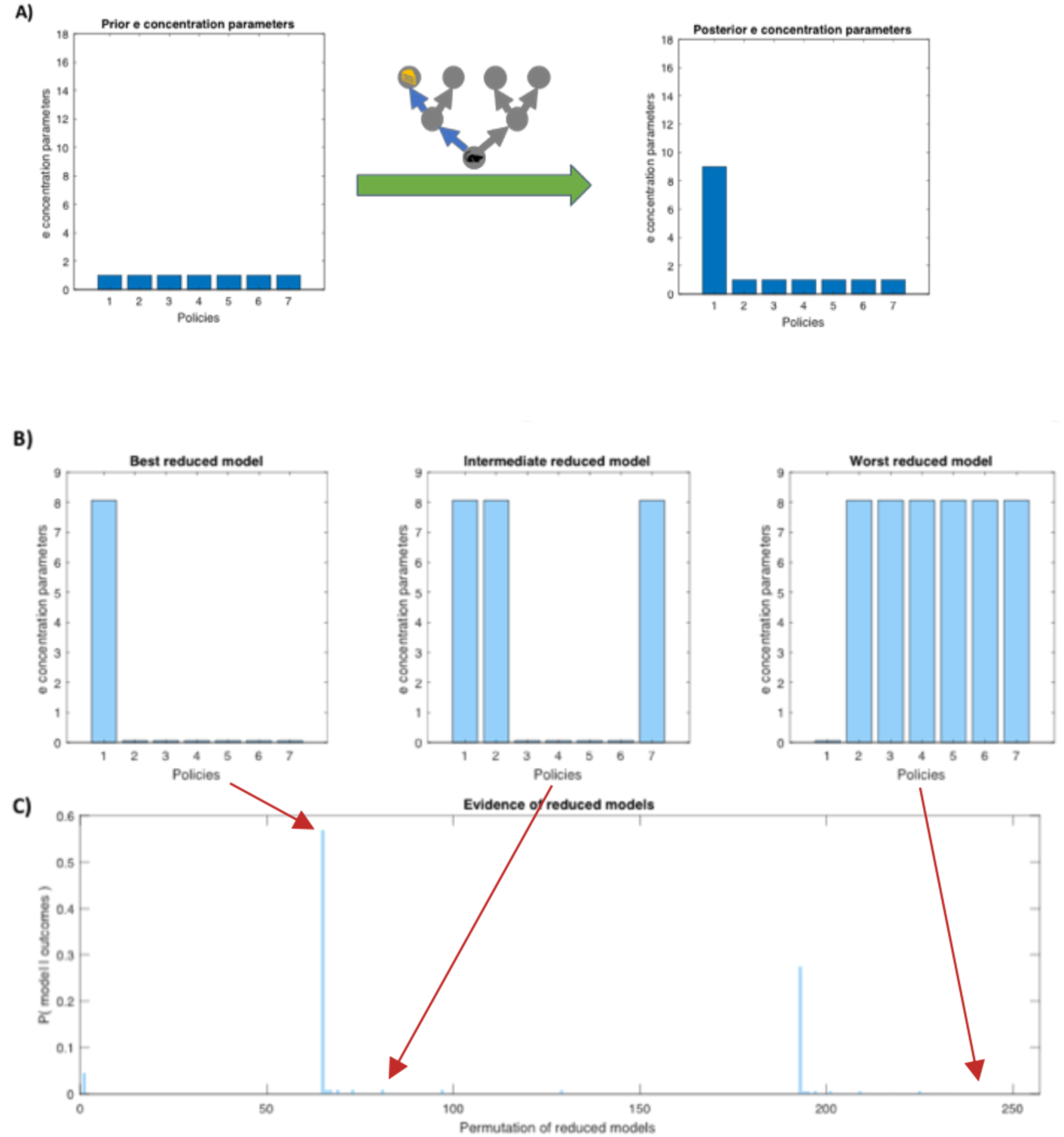
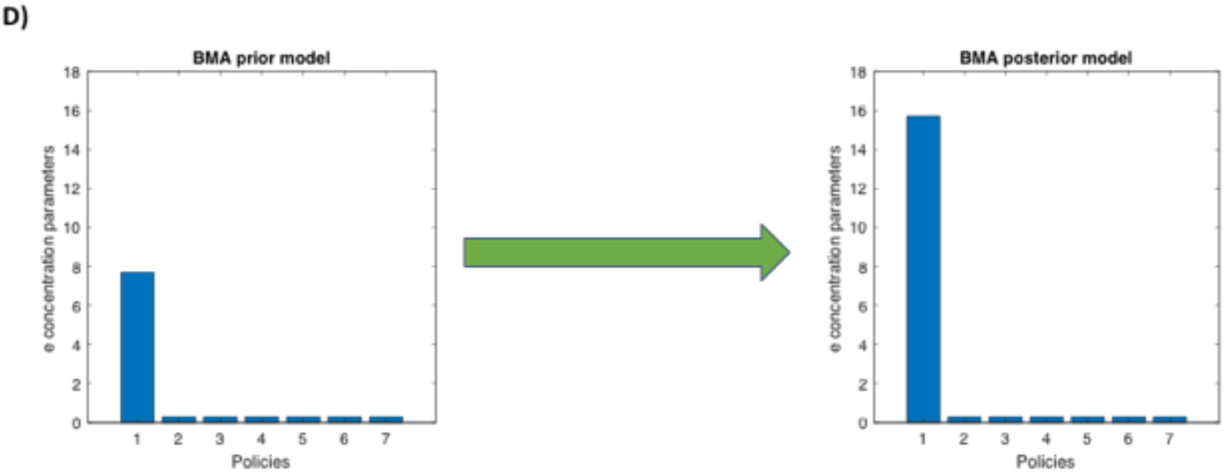
Bayesian Model Averaging (BMA). **(a)** The effect of training on the *e* concentration parameters. The agent pursues policy 1 eight times during the day, and subsequently the *e* parameter for its policy 1 incremented from 1 to 9. **(b)** Example of reduced models. In our case, reduced models are prior *e* concentration parameters that try to better the posterior e concentration observed at the end of the previous day (i.e. part A, right). **(c)** Examples of model evidence. We see the reduced (prior) model increased *e* concentration for policy 1, and decreased concentration for all other policies received the highest model evidence (i.e. it is the best reduced model), whereas models that do the opposite have low model evidence. **(d)** Updating the prior *e* concentration after BMA. The agent first computes the BMA-ed prior *e* concentration (left bar graph), then adds on the amount of learning done during the day to computed the BMA-ed *posterior* e concentration, which is used as the prior for the next day.

The reduced models (Fig A1b) are constructed via strengthening certain policies (increasing their *e* parameters, akin to synaptic strengthening) and weakening others (decreasing *e* parameters, akin to synaptic pruning). The point is to construct many reduced models such that the model space is more likely to contain many good models, and a search through them will pick up those good models (hypothetically, the reduced model space can be arbitrarily large). In our case, we increment the *e* parameter of the to-be-strengthened policies by 8 and divide the *e* of to-be-weakened policies by 2 or 4. The reason for this numerical manipulation is twofold. Firstly, it is more neurobiologically plausible to weaken policies (e.g. via weakening synaptic connections, or in our case, decreasing the *e* parameter by dividing) over time as supposed to “deleting” policies altogether when they are not used. In practice, when the probability of a policy becomes sufficiently small, we can associate this with the pruning of the synapses. Secondly, it is beneficial to construct a large reduced model space, which helps Bayesian model reduction to find a more optimal reduced model. In total, each time model reduction occurs, it iterates through all combinations of reduced policies (since we have 7 policies and we can either strengthen or weaken each one, we have 2^7^= 128 combinations) with the two levels of pruning discussed above for a total of 256 reduced models to average over. Figure A1b, left is an example of a reduced model, in which policy 1 is strengthened (more probable), and all other policies weakened. This is the reduced model with the best model evidence, since it corresponds with the agent’s action during the day (Fig A1a, right).

Now that the probability of each model within the reduced model space is computed (Equation A7, visualized in Fig A1c), we perform Bayesian model averaging get a weighted sum over all the models (Equation A8). The resulting prior (*e_BMA_*) is the optimal set of prior parameters that the agent could have started the previous day with, given the reduced models considered. Finally, the amount of learning (i.e. increases in *e* for policy 1 by 8) is added to this “optimised prior” to get the most optimal posterior *e* concentration, (Fig A1d, right), which is used as the prior concentration for the subsequent day. This is the posterior that the mouse would have reached, had it started with the best prior. This process repeats after each day of training, where the agent continually optimises its parameters to inform better future policy selection.

### Appendix B: Software note

The simulation is constructed using MATLAB (https://www.mathworks.com/products/matlab.html) and the SPM12 software package (https://www.fil.ion.ucl.ac.uk/spm/). Specifically, the DEM toolbox in SPM12 is used to run the Active Inference simulations. All of the scripts used specifically for this experiment can be found on my personal GitHub (https://github.com/im-ant/ActiveInference_PolicyLearning).

